# Inhibitory effect of EGCG-AgNPs and their lysozyme bioconjugates on biofilm formation and cytotoxicity

**DOI:** 10.1101/2022.02.23.481602

**Authors:** Brahmaiah Meesaragandla, Shahar Hayet, Tamir Fine, Una Janke, Liraz Chai, Mihaela Delcea

**Author notes:** **Corresponding Authors**, Liraz Chai, Mihaela Delcea.

## Abstract

Biofilms are multicellular communities of microbial cells that grow on natural and synthetic surfaces. They have become the major cause for hospital-acquired infections because once they form, they are very difficult to eradicate. Nanotechnology offers a new approach to fight biofilm-associated infections. Here, we report on the synthesis of silver nanoparticles (AgNPs) with antibacterial ligand epigallocatechin gallate (EGCG) and the formation of lysozyme protein corona on AgNPs as shown by UV-Vis, dynamic light scattering, and circular dichroism analyses. We further tested the activity of EGCG-AgNPs and their lysozyme bioconjugates on the viability of *Bacillus subtilis* cells and biofilm formation. Our results showed that, although EGCG-AgNPs presented no antibacterial activity on planktonic *Bacillus subtilis* cells, they inhibited *B. subtilis* biofilm formation at concentrations larger than 40 nM and EGCG-AgNP-lysozyme bioconjugates inhibited biofilms at concentrations above 80 nM. Cytotoxicity assays performed with human cells showed a reverse trend, where EGCG-AgNPs barely affected human cell viability, while EGCG-AgNP-lysozyme bioconjugates severely hampered viability. Our results therefore demonstrate that EGCG-AgNPs may be used as non-cytotoxic antibiofilm agents.

## INTRODUCTION

Biofilms are colonies of bacterial cells that form on surfaces and interfaces. Biofilms may be beneficial, for example when they develop on plant roots and protect them from pathogens. Detrimental biofilms clog water and oil pipes, and they may lead to death if they infect catheters at hospitals. Cells in a biofilm are held together by an extracellular matrix (ECM) of proteins, polysaccharides and nucleic acids, which also provides biofilms with mechanical stability and increased antibiotic resistance relative to single cells.^1–2^ In general, antibiotics affect single cells by intervening with cellular growth and proliferation mechanisms, such as cell wall-, nucleic acids- or protein-synthesis.^3–4^ However, bacterial cells in biofilms are less susceptible to antibiotics compared with free-living cells, partly due to the presence of a protective ECM.^5–6^ Therefore, targeting the ECM formation and/or assembly rather than interfering with cellular proliferation, becomes an appealing strategy for biofilm prevention and treatment^7–10^.

Current approaches to prevent biofilm formation include modification of surface topography and surface coatings that detain the cells from sticking to a surface.^11–14^ However, these are temporary treatments because most coatings are quickly covered with self-produced extracellular matrix (ECM) polymers, shielding the anti-fouling coating and allowing the bacterial cells to stick to the surface, despite the coating. Another method related with biofilm eradication includes the use of biocidal molecules, among them silver nanoparticles (AgNPs).^15–16^ AgNPs comprise one of the most predominant nanomaterials in various products, such as fabrics, bandages, deodorizers, food containers and disinfectants.^17–20^ AgNPs are also incorporated into hydrogels, creating hybrid antibacterial wound dressings.^21–24^ These applications exploit AgNPs’ excellent optoelectronic properties (surface plasmon resonance), small size, high surface-to-volume ratio and cost effectiveness.^25–26^ Furthermore, nanosilver is comparatively less reactive than silver ions and is suitable for clinical and therapeutic applications.^27–28^ AgNPs exhibit a broad spectrum of antibacterial and antifungal properties, depending on their shape, size and surface chemistry.^29–37^ The antibacterial activity of AgNPs is related with the release of Ag^+^ ions that may lead to the formation of radical species which damage cells to a lethal extent.^38^

Coating the AgNPs with cationic ligands and other antimicrobial molecules can further improve their antimicrobial activity due to synergistic effects.^39–40^ For example, polyhexamethylene biguanide (PHMB) functionalized AgNPs have shown higher bacteriostatic and bactericidal activity on *Escherichia coli* due to the combined antibacterial effect of AgNPs and PHMB.^41^ Here, we tested the effect of coated AgNPs with epigallocatechin gallate (EGCG) and their lysozyme bioconjugates on biofilm formation. EGCG is a dominant catechin component present in green tea that exhibits various therapeutic properties, including anti-obesity, antiinflammatory, antidiabetic, antitumor effects and antibacterial activity.^42–45^ EGCG is a strong antioxidant, reported to exhibit anti-cancer activity against various cancers (e.g., brain, prostate, pancreatic, and bladder) by inducing apoptosis.^46–53^ Additionally, green tea catechins, including EGCG, can kill bacterial cells.^54–55^ It has been proposed that the bactericidal action of EGCG is due to its ability to generate hydrogen peroxide (H_2_O_2_), which damages bacterial membranes.^42, 56^ On oral administration of EGCG, the rate of ingestion is less than 5% in humans and below 1% in rats.^57–59^ Such low EGCG bioavailability confines its bioactivity *in vivo.* One way to improve the adsorption of EGCG is to adsorb EGCG molecules on AgNPs, which provide large adsorbent surface per unit mass due to their small size.

As drug delivery agents, nanoparticles can be administered in many ways, including oral, nasal, intraocular and parenteral. After coming in contact with biological media, nanoparticles can interact with proteins forming a so-called protein corona and may alter protein structure and its activity.^60–62^ Among the proteins in the body, lysozyme is an antibacterial enzyme which catalyzes the hydrolysis of β1,4-glucosidic linkages between N-acetylglucosamine (NAG) and N-acetylmuramic acid (NAM) in peptidoglycan of the cell wall, particularly in Gram-positive bacteria.^63^ Lysozyme is a small, monomeric, and globular protein, abundant in various body fluids including serum (4-13 mg/L), saliva, tears and human milk.^64–66^ It consists of 130 amino acid residues with a molecular mass of 14.7 kDa and belongs to the α+β class of proteins. The compact structure of lysozyme is stabilized by four disulfide bonds and its surface is typically polar, whereas its inner part is almost hydrophobic. It has been shown that Gram-positive bacteria are more susceptible to the antibacterial activity of lysozyme than Gram-negative bacteria.^67–68^

In the present study, we have synthesized EGCG-AgNPs to study their antibacterial activity against the Gram-positive wild type bacterium *Bacillus subtilis* through bacterial growth and biofilm formation assays. To enhance the antibacterial activity of the AgNPs, we also conjugated EGCG-AgNPs with the antibacterial lysozyme (EGCG-AgNP-lysozyme). Indeed, it has been shown that decorating AgNPs with lysozyme and other bactericides may enhance their antibacterial spectrum.^39,40,69^ To attest for possible application of EGCG-AgNPs and their lysozyme bioconjugates in the human body, we also studied their cytotoxicity effect against human cells (see scheme 1 for an overview of the study). This study highlights the design of safe and effective EGCG-AgNPs antibiofilm agents.

**Scheme 1.**
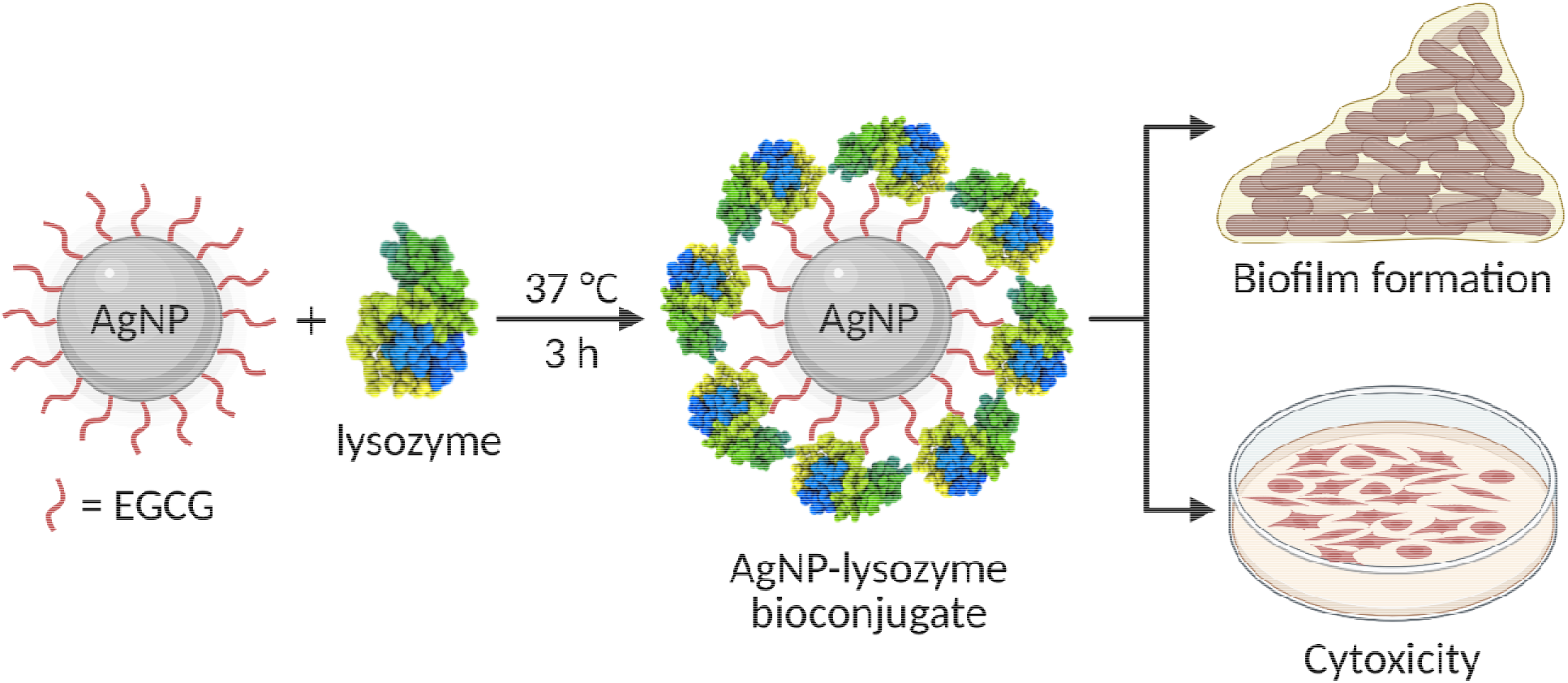
Overview of this study illustrating functionalization of AgNPs with EGCG and the formation of a bioconjugate with lysozyme. We have studied the inhibitory effect of EGCG-AgNPs and their lysozyme bioconjugates on biofilm formation and also their cytotoxicity.

## RESULTS AND DISCUSSION

### Synthesis and characterization of EGCG-AgNPs

AgNPs were prepared by the reduction of Ag^+^ in a solution of silver nitrate (AgNO_3_) in the presence of EGCG, a polyphenol used as reducing agent as well as an NP stabilizing agent due to the availability of hydroxyl (-OH) groups.^70–71^ The formation of AgNPs was confirmed by the occurrence of a UV-Vis absorption band, peaking at 413 nm, which is characteristic of surface plasmon resonance (SPR) (Figure 1A). TEM analysis showed spherical EGCG-AgNPs with a diameter of 14 ± 5 nm (Figure 1B), whereas DLS showed a hydrodynamic diameter (d_H_) of 36 ± 2 nm and 130 ± 10 nm (Figure 1C). The larger d_H_ values of the AgNPs, obtained with DLS, compared to those observed in the TEM images, may be attributed to the formation of EGCG-AgNPs dimers (~ 36 nm), or aggregates (~ 130 nm). NP aggregates may form either due to surface-aggregation processes as a result of poor coverage or weak binding of the ligand, or due to aggregation of the ligand itself at the surface of the NPs. The former would affect the SPR and result with a red shift in the absorbance, however, the relatively sharp absorbance band of the EGCG-AgNPs and the lack of an additional, red-shifted peak, implies that the population of larger NPs originates from surface-aggregation of EGCG ligands rather than from aggregation of NPs. Figure S1A and S1B show the time-dependent UV-Vis spectra of EGCG-AgNPs in water and in PBS (pH 6.2), respectively. EGCG-AgNPs were stable in water, as indicated by the insignificant changes in the absorption band peak position and intensity in the course of 24 hours (Figure S1A).

**Figure 1.**
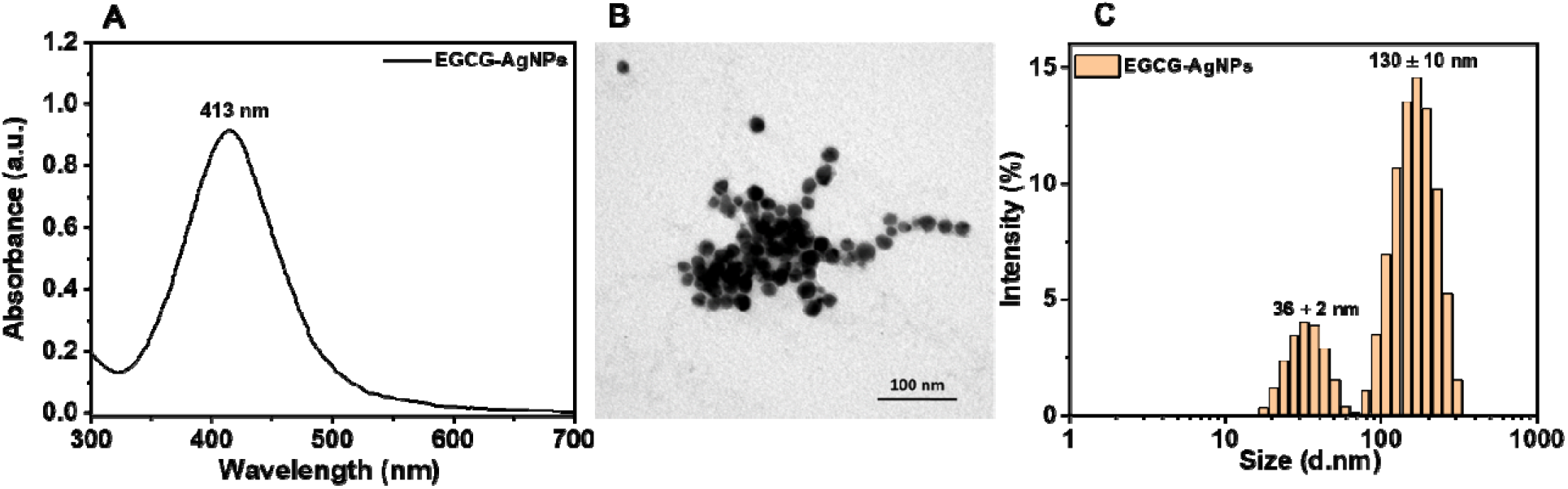
Characterization of EGCG-AgNPs. UV-Visible spectrum (A), TEM image (B) and size distribution (C) of EGCG-AgNPs in water.

In contrast, adding the EGCG-AgNPs into PBS buffer, resulted in a blue shift and a decrease in the intensity of the SPR absorption band (Figure S1B), indicating that AgNPs are unstable in PBS buffer.^72–73^ The reduced stability of EGCG-AgNPs in PBS relative to water may be explained by partial substitution of EGCG with salt and it stands in agreement with previous reports of the aggregation of AgNPs upon the addition of salt.^74^ This was further supported by increased negative zeta potential values of EGCG-AgNPs in PBS (−35 mV) when compared to water (−27 mV) (Figure S2).

### Interaction of human lysozyme with EGCG-AgNPs

The interaction of lysozyme with EGCG-AgNPs in PBS (pH 6.2) was characterized by UV-Vis, DLS, and zeta potential analyses. We fixed the concentration of lysozyme (6.8 μM), and added EGCG-AgNPs at increasing final concentrations. Addition of EGCG-AgNPs to a lysozyme/PBS buffer solution resulted in a red shift of the absorption band (Figure 2A). Interestingly, the peak position and the intensity of the SPR band remained unchanged even after incubation at 37 □ for 3 h. The red shift of the SPR absorption band is indicative of strong interactions between EGCG-AgNPs and lysozyme molecules, and it is also supported by an increase in the overall d_H_ for EGCG-AgNP-lysozyme bioconjugates (Figure 2B). However, the intensity of the SPR peak remained unchanged, suggesting that despite the increased size of the AgNP-lysozyme bioconjugates relative to EGCG-AgNPs, the AgNPs remained stable in solution. A possible explanation would be steric repulsion between lysozyme molecules. Increasing the lysozyme concentration (6.8, 20.4, 34 μM) while keeping the EGCG-AgNPs concentration fixed (100 nM) did not affect the SPR absorption peak (Figure S3), indicating that a minimum lysozyme concentration (6.8 μM) is sufficient to form stable EGCG-AgNP-lysozyme bioconjugates, possibly due to saturation of the binding sites at the AgNPs surface. In fact, measurement of the concentrations of lysozyme that was bound to EGCG-NPs and lysozyme that remained free in solution revealed that ~20% of the lysozyme was bound to the NPs while 80% was unbound (see experimental section and Figure S4).

**Figure 2.**
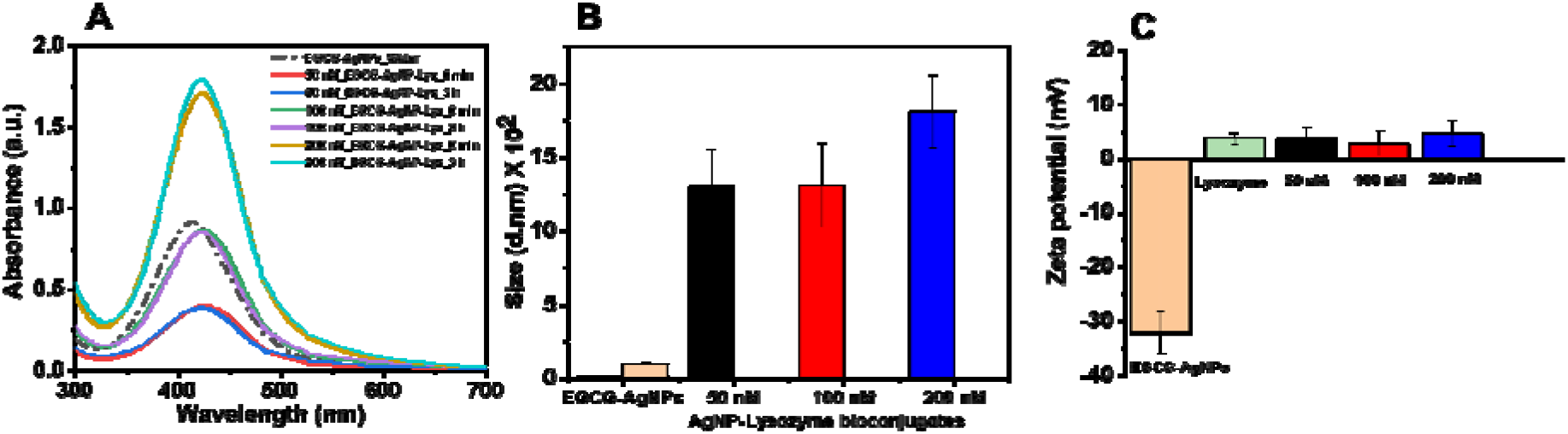
Characterization of EGCG-AgNP-lysozyme bioconjugates. (A) UV-Visible spectra of EGCG-AgNP-lysozyme bioconjugates before and after 3 h incubation in PBS buffer (pH 6.2) at 37 □. (B) Size (d_H_) of EGCG-AgNPs and their AgNP-lysozyme bioconjugates after 3 h incubation. (C) Zeta potential of EGCG-AgNPs, lysozyme, and their corresponding AgNP-lysozyme bioconjugates at pH 6.2. Lysozyme concentration was fixed to 6.8 μM and AgNPs concentration was varied (50, 100, 200 nM).

Interaction of EGCG-AgNPs with lysozyme was characterized by a positive surface charge, in agreement with the isoelectric point of lysozyme (10.7)^64–66^ being larger than the working pH (6.2). The positive zeta potential of the AgNPs confirmed lysozyme adsorption onto the EGCG-AgNPs and formation of EGCG-AgNP-lysozyme bioconjugates (Figure 2C). Importantly, despite the lower zeta potential (absolute value) of the EGCG-AgNP-lysozyme bioconjugates relative to EGCG-AgNPs, the EGCG-AgNP-lysozyme bioconjugates remained stable in solution, confirming further the steric stabilization of the NPs by lysozyme molecules in PBS buffer.

### Effect of EGCG-AgNPs on the secondary structure of lysozyme

Figure 3 shows the CD spectra of lysozyme alone and lysozyme with EGCG-AgNPs following 3h incubation in varying concentrations, ranging between 10 nM-200 nM. The CD spectra of lysozyme at pH 6.2 exhibit two negative minima at 208 nm and 222 nm, which is characteristic of α-helical content, corresponding to π–π* and n–π* transitions of the peptide bonds, respectively.^75^ Interestingly, adding EGCG-AgNPs to lysozyme solution affected the intensity and shape of CD spectra in a concentration-dependent manner. Specifically, increasing the concentration of EGCG-AgNPs resulted with decreased ellipticity and transition of lysozyme structure from α-helix to β-sheet at concentrations ≥ 100 nM. Loss of the characteristic alpha helical peak at 208 nm upon addition of EGCG-AgNPs, is shown in Fig. 3B. Table S1 shows relative fractions of secondary structures of lysozyme, calculated by the Bestsel software by best fitting the CD spectra with linear combinations of spectra of known protein structures.^76^ This analysis confirms the qualitative transition of lysozyme from α-helix-to β-sheet rich, quantifying the reduction in the α-helical fraction of the lysozyme from 20.5 % to 3.4 % in the presence of EGCG-AgNPs (200 nM) after 3 h incubation. Intramolecular hydrogen bonds stabilize lysozyme molecules and lead to an α-helical structure. Hydrogen bonding between hydroxyl-EGCG and lysozyme may interfere with the lysozyme intramolecular interaction and stand responsible for the structural changes of lysozyme.

**Figure 3.**
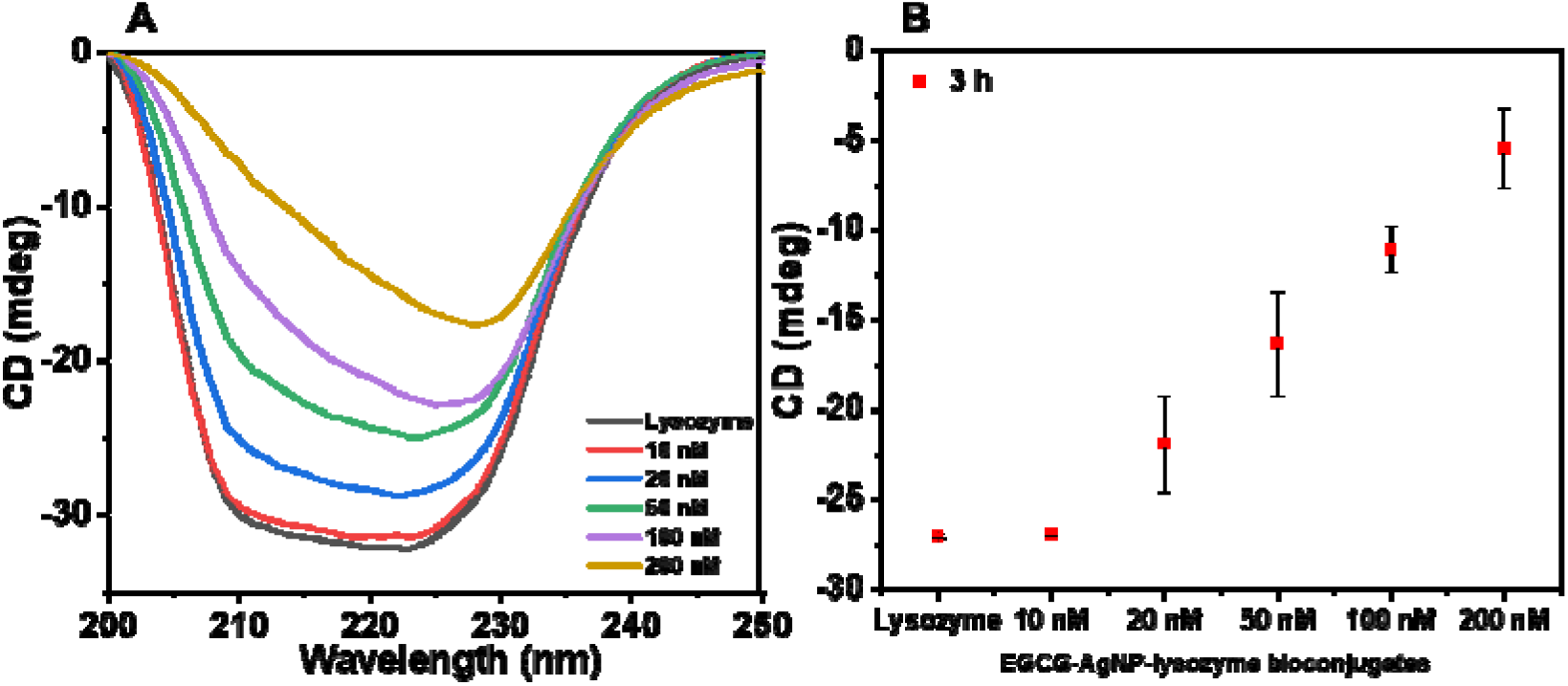
(A) CD spectra of human lysozyme (6.8 μM) and the same concentration in EGCG-AgNP-lysozyme bioconjugates with increased concentrations of AgNPs in PBS buffer (pH 6.2) after 3 h incubation at 37 □. (B) Plot of ellipticity values at 208 nm for lysozyme and EGCG-AgNP-lysozyme bioconjugates at pH 6.2.

### Effect of EGCG-AgNPs and EGCG-AgNP-lysozyme bioconjugates on *Bacillus subtilis* bacterial growth and biofilm formation

We examined the effect of the EGCG-AgNPs and EGCG-AgNP-lysozyme bioconjugates on cell viability and biofilm formation of WT *B. subtilis.* The growth curves were similar in the presence and in the absence of EGCG-AgNPs (Figure 4A, log/semi-log plot shown in Figure S5A), suggesting that EGCG-AgNPs had no antibacterial activity on planktonic *B. subtilis* cells. This result is surprising in light of the antibacterial activity of both free EGCG and free AgNPs.^77^ We speculate that the functional groups of EGCG attach to AgNPs and therefore, EGCG loses its antibiotic activity. At the same time, EGCG binding to AgNPs prevents the leaching of Ag^+^ ions, which in turn compromises nanoparticle antibiotic activity.

**Figure 4.**
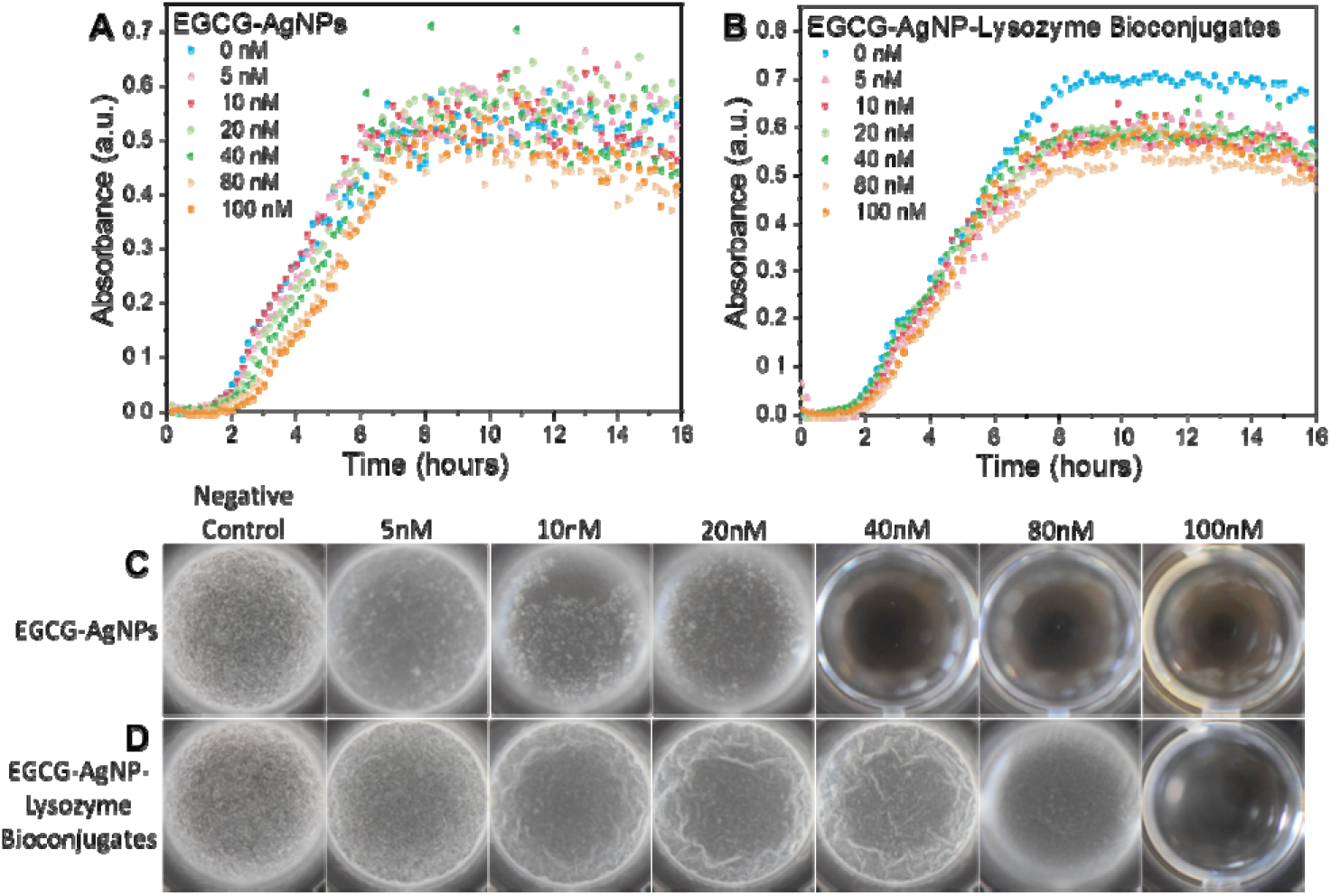
Effect of EGCG-AgNPs and their lysozyme bioconjugates on WT *B. subtilis* growth curves and biofilm assays. Growth curves in the presence of (A) EGCG-AgNPs and (B) EGCG-AgNP-lysozyme bioconjugates in PBS at pH 6.2. The concentration of the NPs is specified in the legends. Biofilm formation by *B. subtilis* in the absence and in the presence of increasing concentrations of EGCG-AgNPs (C) and EGCG-AgNP-lysozyme bioconjugates (D).

In contrast to the compromised antibacterial activity against cells in cultures, EGCG-AgNPs inhibited *B. subtilis* in biofilm forming conditions at concentrations larger than 40 nM. Inhibition of biofilm formation could be attributed to a biocidal effect of EGCG-AgNPs, however, the growth curves in figure 4A rule out this possibility, because they remained unchanged even in the presence of EGCG-AgNPs at a 100 nM concentration. Therefore, EGCG-AgNPs may have interfered with biofilm formation pathways, such as the expression of ECM components or prevention of their proper assembly into a network^78^, as well as communication by quorum sensing.^79^

Similar to EGCG-AgNPs, when these NPs were additionally functionalized with lysozyme, they showed no biocidal activity on *B. subtilis* cells in the concentration range we have used (Figure 4B, log/semi-log plot shown in Figure S5B). However, the lysozyme corona reduced the effect of EGCG-AgNPs on *B. subtilis* biofilms. Specifically, biofilm inhibition occurred at concentrations larger than 80 nM EGCG-AgNP-lysozyme bioconjugates, which is larger than the 40 nM inhibitory concentration for biofilm inhibition by EGCG-AgNPs. The decrease in biofilm inhibition by the EGCG-AgNP-lysozyme bioconjugates may be either due to the blockage of negatively charged EGCG moieties by the lysozyme or due to the loss of lytic activity of lysozyme against bacterial cell wall when it is bound to EGCG-AgNPs.

### Cytotoxicity of EGCG-AgNPs and EGCG-AgNP-lysozyme bioconjugates

We have further investigated the toxicity of EGCG-AgNPs and EGCG-AgNP-lysozyme bioconjugates on HUVEC cells as model system for endothelial cells, which are forming the inner layer of blood vessels. Figure 5 shows the HUVEC cell viability data after treatment with different concentrations of EGCG-AgNPs, EGCG-AgNP-lysozyme bioconjugates. EGCG-AgNPs are barely cytotoxic below 200 nM, however, a decrease in cell viability was observed at 200 nM EGCG-AgNPs. EGCG-AgNPs present a lower cytotoxicity when compared to EGCG-AgNP-lysozyme bioconjugates, the latter showing decreased cell viability already at 10 nM. The differences between the cytotoxicity of EGCG-AgNPs and lysozyme-functionalized EGCG-AgNPs may be related with faster uptake rates of the lysozyme bioconjugates by the cells. An additional possibility would be that human cells are more affected by structural changes of lysozyme, in particular of β-sheet structures, relative to bacterial cells.

**Figure 5.**
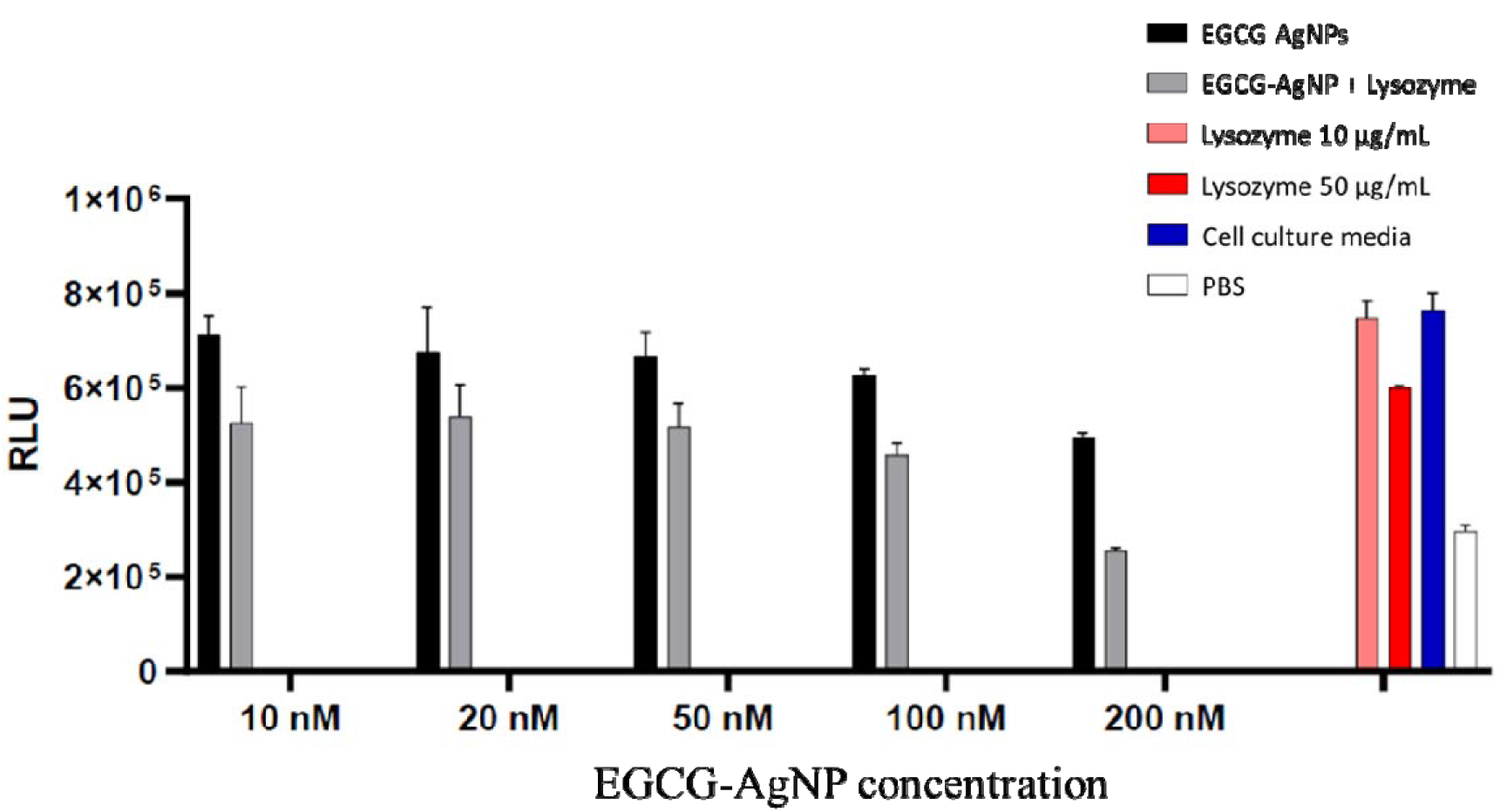
Cytotoxicity assay of HUVEC cells for different concentrated EGCG-AgNPs (10, 20, 50, 100, 200 nM) and the corresponding AgNP-lysozyme bioconjugates in endothelial cell culture media supplemented with fetal bovine serum (FBS). Measurement data was normalized as explained in the methods section.

## CONCLUSIONS

We have synthesized EGCG-AgNPs and studied their interaction with lysozyme. Conjugation of EGCG-AgNPs with lysozyme increased the stability of these NPs in PBS and it induced structural changes of lysozyme from a dominant α-helical to dominant β-sheet fraction in a concentration dependent manner.

Studies of the inhibitory effect of EGCG-AgNPs and lysozyme-conjugated EGCG-AgNPs on Gram-positive wild type *B. subtilis* biofilms showed that EGCG-AgNPs inhibited the formation of *B. subtilis* biofilm above 40 nM, whereas the lysozyme bioconjugates of EGCG-AgNPs were less effective and inhibited biofilms at higher concentration (≥ 80 nM). Interestingly, EGCG-AgNPs showed lower cytotoxicity when compared to EGCG-AgNP-lysozyme bioconjugates. Strikingly, both EGCG-AgNPs and lysozyme conjugated EGCG-AgNPs did not affect the viability of bacterial cells, suggesting that (i) bound EGCG is not harmful to bacterial cells, and (ii) NPs need to be uptaken into the cells in order to cause damage.

Our results demonstrate that EGCG-AgNPs can be used as an antibiofilm agent against *B. subtilis* as they showed lower toxicity against human cells and significant inhibitory effect on biofilm formation.

## EXPERIMENTAL SECTION

### Materials

Human lysozyme, (-)-Epigallocatechin gallate (EGCG), phosphate buffer saline (PBS) and silver nitrate (AgNO_3_) were purchased from Sigma-Aldrich (Taufkirchen, Germany). EtOH, HCl, and NaOH were purchased from Roth (Karlsruhe, Germany). The concentration of lysozyme was measured spectrophotometrically using a molar extinction coefficient of 38940 mol^-1^cm^-1^ at 280 nm. The water used was purified through an ultrapure water system, Millipore system Sartorius Stedim Biotech (Göttingen, Germany).

PBS was prepared by dissolving 0.137 M NaCl, 0.0027 M KCl, 0.01 M Na_2_HPO_4_, 0.0018 M KH_2_PO_4_ in triple-distilled water and adjusting for pH with HCl. Except NaCl, all chemicals were purchased from Sigma-Aldrich (Darmstadt, Germany). NaCl was purchased from J.T. Baker-Avantor (Center Valley, PA, USA). The water was purified through Barnstead GenPure water purification system (Thermo Scientific).

### Epigallocatechin gallate (EGCG)-AgNPs

EGCG (10 mM) solution was prepared by dissolving 4.58 mg of EGCG in 1 mL deionized water which was pre-adjusted to pH 8, then it was added dropwise to 30 mL of 1 mM AgNO_3_ (5.03 mg in 30 mL water) solution with continuous stirring at 27 □ for 2 h. AgNO_3_ solution turned from colorless to yellowish brown upon addition of EGCG solution.

### EGCG-AgNP-lysozyme interaction

EGCG-AgNPs were added to lysozyme in PBS (pH 6.2) and the mixture was incubated at 37 □ for 3 h prior to characterization. For the analysis of protein structural changes, a fixed concentration of lysozyme (100 μg mL^-1^) was mixed with various concentrations of EGCG-AgNPs (10, 20, 50, 100, 200 nM) and then incubated at 37 □ for 3 h. Similarly, various concentrations of lysozyme (100, 300 and 500 μg/mL) were used for UV-Vis analyses.

### EGCG-AgNP-lysozyme bioconjugates

EGCG-AgNPs (1 μM stock solution) were added in PBS buffer (pH 6.2) containing 100 μg/mL lysozyme and incubated at 37 °C for 4 h. EGCG-AgNP-lysozyme bioconjugates were obtained by centrifuging the above mixture at 17,000 g at room temperature for 30 min and resuspending in PBS buffer. The bioconjugates were diluted with PBS buffer to various concentrations (5, 10, 20, 50, 80, 100 nM).

### Determination of lysozyme concentration on the AgNPs surface

Protein concentration on the EGCG-NPs and the unbound protein that remained in solution was determined by measuring absorbance at 280 nm (A_280_) using a DeNovix DS-11 FX+ spectrophotometer. A lysozyme calibration curve was prepared by measuring the A_280_ of a series of lysozyme dilutions of known concentrations (1mg/ml – 0.0156 mg/ml). Protein solution of lysozyme (100 μl of 68 μM) was mixed with aqueous dispersion of EGCG-AgNPs (100 nM) to a final concentration of 6.8 μM. The EGCG-Ag-lysozyme mixture was incubated for 3 h at 37 °C and the EGCG-NP-lysozyme bioconjugates were precipated by centrifugation (17g X 30 min). The unbound (supernatant) protein was removed from the AgNP-lysozyme mixture and the EGCG-AgNP-lysozyme bioconjugates pellet was resuspended in PBS. Absorbance of the supernatant and pellet was measured and the lysozyme concentration was determined by plugging-in the background subtracted absorbance in the absorbance versus concentration calibration curve (see Figure S4 for the calibration curve and lysozyme concentration determination).

### Circular dichroism spectroscopy measurements

The far-UV circular dichroism (CD) spectra of lysozyme after incubation with AgNPs were recorded on a Chirascan spectrophotometer (Applied Photophysics, Leatherhead, UK). The spectrophotometer was purged with nitrogen gas before the experiments. Measurements were scanned between 200 and 250 nm with an average of 5 scans using a 5 mm path length cuvette (Hellma Analytics, Müllheim, Germany). All spectra were measured at room temperature with a band width of 1.0 nm. The final data was obtained by subtracting the buffer contribution from the original protein spectra. The fractional contents of the secondary structures were calculated by the Bestsel software.^75^

### UV-Vis absorption spectroscopy

DeNovix DS-11 FX+ spectrophotometer (Biozym Scientific GmbH, Germany) was used to obtain the absorption spectra for EGCG-AgNPs, and the same after interacting with lysozyme. All the samples were measured between 200 to 850 nm using 10 mm path length cuvette at room temperature.

### Dynamic light scattering (DLS) and zeta potential measurements

Hydrodynamic diameter (d_H_) and zeta potential for EGCG-AgNPs and the corresponding lysozyme bioconjugates were determined using Zetasizer Ultra (Malvern Instruments, Kassel, Germany). Except lysozyme bioconjugates, all the samples were prepared as described above and filtered through a 0.2 μm filter. Before measurements, samples were equilibrated for 10 min at room temperature. Each size measurement was recorded allowing 20 runs per measurement with a run duration of 5 s. An average of five separate measurements was used to assess the size of the samples. DTS1070 cells were used to measure the zeta potential of AgNPs and their lysozyme bioconjugates. A voltage of 180/40 V was used for the samples measured in deionized distilled water and PBS, respectively. All measurements were acquired at room temperature with an equilibration time of 5 min (20 runs per each measurement) between each measurement. The reported zeta potential is an average of five independent measurements.

### Transmission Electron Microscopy

The flotation method was used for the negative staining procedure. AgNPs were allowed to adsorb onto a glow-discharged Pioloform carbon-coated 400-mesh grid for 5 min. The grid was then transferred onto two droplets of deionized water and finally onto a drop of 1 % aqueous uranyl acetate for 30 s. After blotting with filter paper and airdrying, the samples were examined with a transmission electron microscope LEO 906 (Carl Zeiss Microscopy Deutschland GmbH, Oberkochen, Germany) at an acceleration voltage of 80 kV. For image acquisition, a wide-angle dual speed CCD camera Sharpeye (Tröndle, Moorenweis, Germany) was used, operated by the ImageSP software. All micrographs were edited by using Adobe Photoshop CS6.

### Biofilm and Growth Curve Assays

Liquid culture of WT *B. subtilis* was grown in LB broth at 37 °C for 16 h. For the biofilm formation assay, 2 μL of starting culture were added to 88 μL of MSgg medium in a 96-well plate and incubated (24 h, 30 °C) with 10 μL EGCG-AgNPs or with EGCG-AgNP-lysozyme bioconjugates to achieve various final concentrations: 5, 10, 20, 40, 80, and 100 nM. Biofilms were captured with a Nikon D3300 camera with a micro Nikkor 85 mm lense. For the growth curve assay, 2 μL of starting culture were added to 88 μL of LB broth in a 96-well plate with 10 μL EGCG-AgNP-Lysozyme bioconjugates in various final concentrations, as specified above for biofilms. Bacterial growth was measured in a microplate reader (Tecan Spark 10M, Tecan Trading AG, Switzerland) for 16 h at 37 °C with 180 RPM orbital shaking.

EGCG-AgNPs stock solutions were prepared in triple-distilled water, and EGCG-AgNP-lysozyme stock solutions were prepared in PBS buffer. Control experiments without EGCG-AgNPs were performed with water as a substitute of EGCG-AgNPs and PBS buffer as substitute for EGCG-AgNP-lysozyme bioconjugates.

### Cell Viability Assay

Cell viability assays were carried out following manufacturer’s instructions from the CellTiter-Glo 2.0 assay from Promega (Madison, USA). In brief, 5×10^4^ human umbilical vein endothelial cells (HUVEC)/mL were seeded in endothelial cell growth medium MV (Promocell, Heidelberg, Germany) with SupplementMix. Cells were incubated for 3 h in an opaque 96-well plate at 37 °C and 5% CO_2_. Then, media was removed and different concentrations of AgNPs were added, indicated in the respective graphs. After 24 h incubation at 37°C and 5% CO_2_, 100 μL of CellTiter-Glo 2.0 was added to each well and luminescence signal was measured in a Cytation 5 imaging reader (BioTek instruments Inc., Winooske, USA) after 10 min incubation.

## ASSOCIATED CONTENT

### Supporting Information

UV-Vis absorption spectra and Zeta potential of EGCG-AgNPs in water and PBS; Zeta potential of EGCG-AgNPs in water and PBS; UV-Visible spectra of EGCG-AgNP-lysozyme bioconjugates; Determination of the EGCG-NPs-bound and unbound lysozyme fraction; Growth curves of WT *B. subtilis* in the presence of EGCG-AgNPs and of EGCG-AgNP-lysozyme bioconjugates; Percentages of secondary structures of lysozyme in presence of EGCG-AgNPs at pH 6.2.

## ACKNOWLEDGMENT

We acknowledge Dr. Rabea Schlüter (University of Greifswald) for TEM analyses. BM and MD acknowledge the German Federal Ministry of Education and Research (BMBF) for the financial support (project NanoImmun FKZ03Z22C51).

