## Supporting information for "Inhibitory effect of EGCG-AgNPs and their lysozyme bioconjugates on biofilm formation and cytotoxicity"

*Corresponding authors:

**Figure S1**. UV-Vis spectra of EGCG-AgNPs in water and PBS.

**Figure S2**. Zeta potential of EGCG-AgNPs in water and PBS.

**Figure S3.** UV-Visible spectra of EGCG-AgNP-lysozyme bioconjugates.

**Figure S4.** Determination of the EGCG-NPs-bound and unbound lysozyme fractions.

**Figure S5.** Growth curves of WT *B. subtilis* in the presence of EGCG-AgNPs and of EGCG-AgNP-lysozyme bioconjugate in log/semi-log plots.

**Table S1**. Percentages of secondary structures of lysozyme in presence of EGCG-AgNPs at pH 6.2.

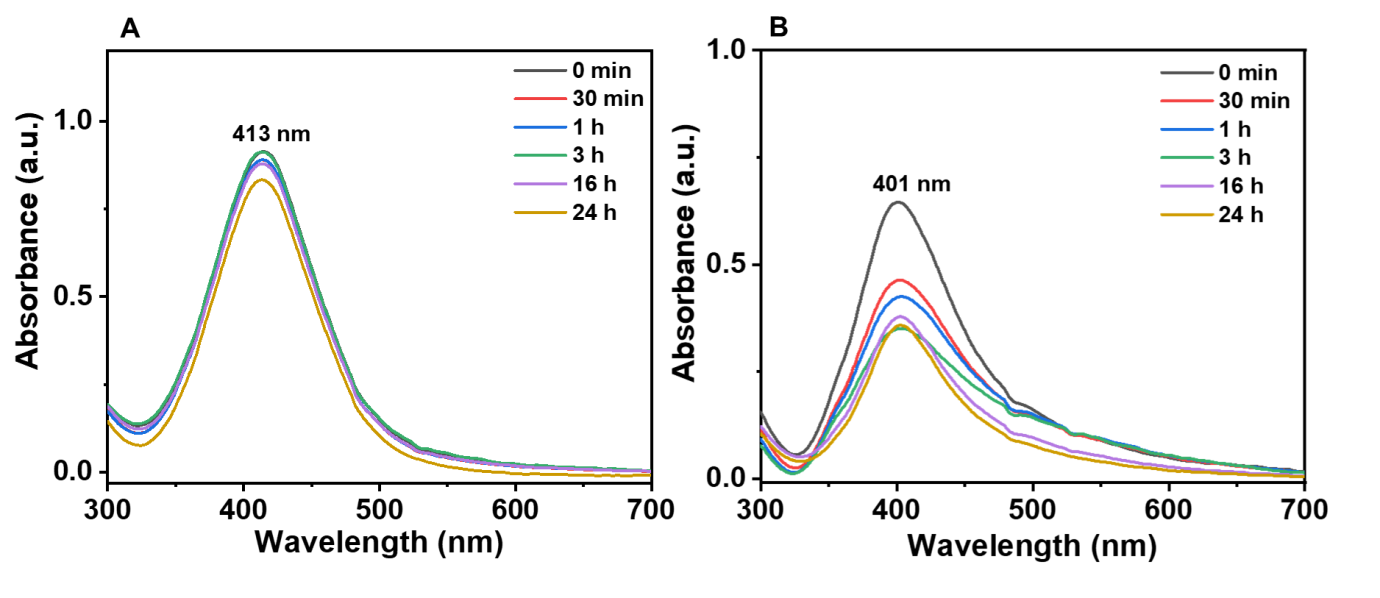

**Figure S1.** UV-Vis spectra of EGCG-AgNPs in water (A) and in PBS buffer at pH=6.2 (B).

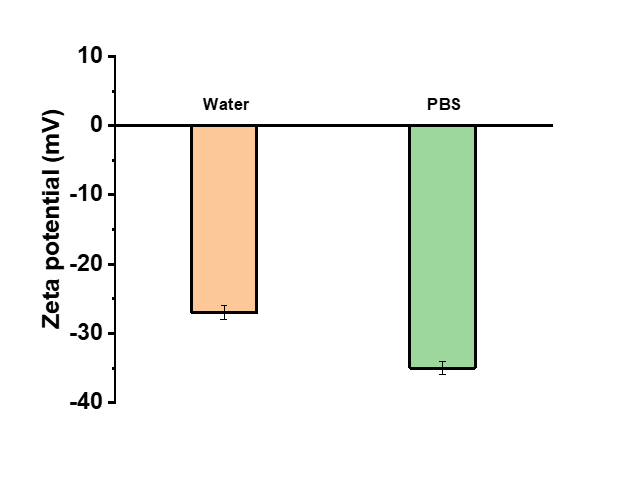

**Figure S2.** Zeta potential of EGCG-AgNPs in water and in PBS at pH 6.2 (samples were measured in PBS after 30 min).

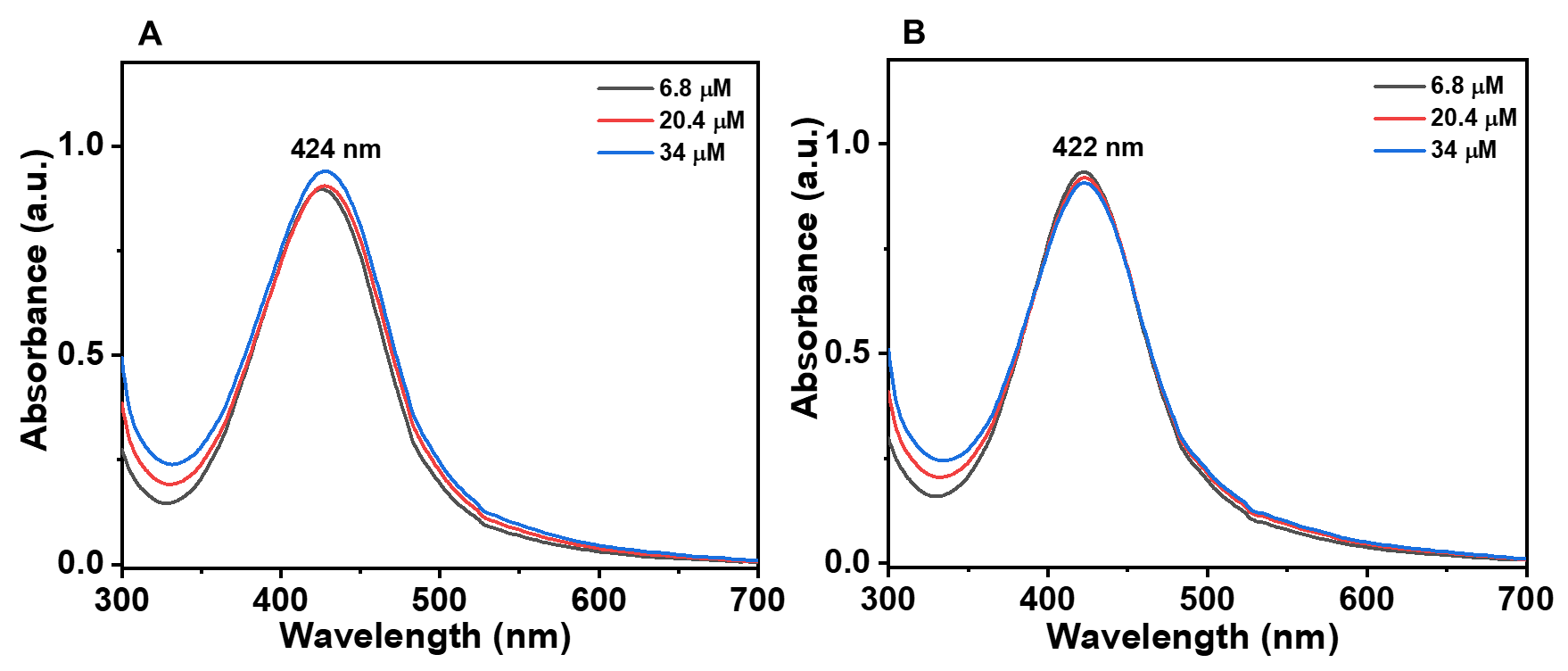

**Figure S3.** UV-Visible spectra of EGCG-AgNP-lysozyme bioconjugates before (A) and after 3 h incubation in PBS buffer (pH 6.2) at 37 °C (B). EGCG-AgNPs concentration was fixed to 100 nM and lysozyme concentration was varied (6.8, 20.4, 34 µM).

A)

| Absorbance (A_280_) | Concentration in mg/mL |
| --- | --- |
| 2.6375 | 1 |
| 1.3135 | 0.5 |
| 0.6328 | 0.25 |
| 0.3181 | 0.125 |
| 0.151 | 0.0625 |
| 0.0771 | 0.03125 |
| 0.044 | 0.0156 |

B)

Standard curve

| concentration in mg/mL | absorbance at 280 nm |
| --- | --- |
| 1 | 2.6375 |
| 0.5 | 1.3135 |
| 0.25 | 0.6328 |
| 0.125 | 0.3181 |
| 0.0625 | 0.151 |
| 0.03125 | 0.0771 |
| 0.0156 | 0.044 |

C)

|  | Absorbance (280 nm) | Concentration (µg/ml) | Concentration (µM) | Adsorbed/unbound fraction |
| --- | --- | --- | --- | --- |
| EGCG-NPs pellet | 0.286 ± 0.006 | 15 ± 6 | 1.0 ± 0.4 | 17 ± 7  17 ± 7 |
| EGCG-NPs supernatent | 0.178 ± 0.008 | 71 ± 7 | 4.9 ± 0.5 | 83 ± 8 |

**Figure S4**. Determination of the lysozyme fraction that was bound to EGCG-NPs following incubation with lysozyme and the remaining unbound fraction. The absorbance of lysozyme in the range 0.0156 – 1 mg/mL was measured at 280 nm (A) and used to plot a calibration curve of absorbance versus concentration was (B). EGCG-NPs bioconjugates were precipitated by centrifugation to separate pellet and sup and the lysozyme concentration was determined in both fractions via absorbance measurements (C). The concentration of the lysozyme solution used during the incubation with EGCG-NPs was 6 µM, as was determined by absorbance measurement.

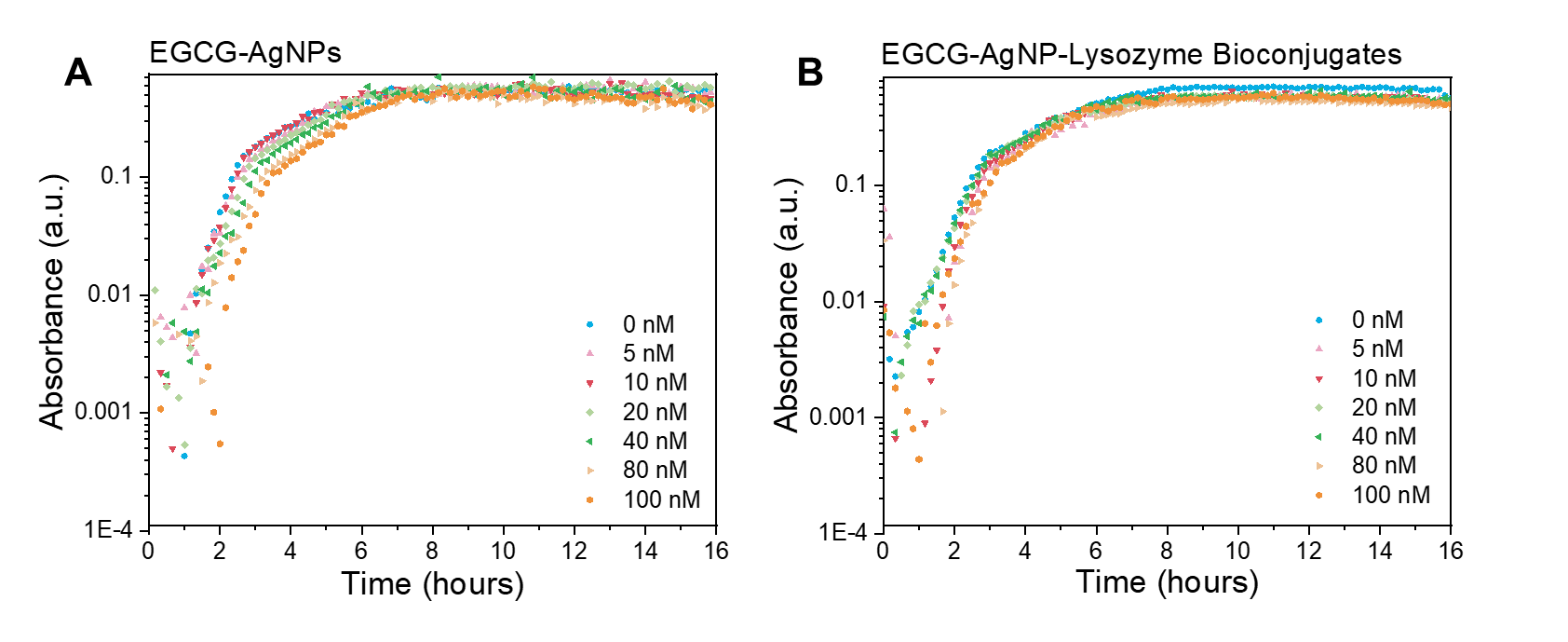

**Figure S5.** Log/semi-log plots of WT *B. subtilis* growth curves in the presence of EGCG-AgNPs (A) and in the presence of EGCG-AgNP-lysozyme bioconjugates in PBS at pH 6.2 (B).

**Table S1.** Percentages of secondary structures of lysozyme and the same upon interaction with different concentrations of EGCG-AgNPs at pH 6.2 after 3h incubation at 37 °C.

| **After incubation (3 h)** | **Helix (%)** | **Antiparallel (%)** | **Parallel (%)** | **Turns (%)** | **Others (%)** |
| --- | --- | --- | --- | --- | --- |
| Lysozyme | 20.5 | 13.5 | 10.4 | 15.6 | 40.0 |
| 10 nM | 20.2 | 14.8 | 10.8 | 15.6 | 38.6 |
| 20 nM | 17.2 | 17.0 | 12.1 | 15.0 | 38.7 |
| 50 nM | 14.1 | 20.8 | 12.4 | 14.7 | 38.1 |
| 100 nM | 9.7 | 22.7 | 13.9 | 14.5 | 39.2 |
| 200 nM | 3.4 | 30.2 | 10.9 | 15.6 | 40.0 |
